# SARS-CoV-2 Lambda Variant Remains Susceptible to Neutralization by mRNA Vaccine-elicited Antibodies and Convalescent Serum

**DOI:** 10.1101/2021.07.02.450959

**Authors:** Takuya Tada, Hao Zhou, Belinda M. Dcosta, Marie I. Samanovic, Mark J. Mulligan, Nathaniel R. Landau

## Abstract

The SARS-CoV-2 lambda variant (lineage C.37) was designated by the World Health Organization as a variant of interest and is currently increasing in prevalence in South American and other countries. The lambda spike protein contains novel mutations within the receptor binding domain (L452Q and F490S) that may contribute to its increased transmissibility and could result in susceptibility to re-infection or a reduction in protection provided by current vaccines. In this study, the infectivity and susceptibility of viruses with the lambda variant spike protein to neutralization by convalescent sera and vaccine-elicited antibodies was tested. Virus with the lambda spike had higher infectivity and was neutralized by convalescent sera and vaccine-elicited antibodies with a relatively minor 2.3-3.3-fold decrease in titer on average. The virus was neutralized by the Regeneron therapeutic monoclonal antibody cocktail with no loss of titer. The results suggest that vaccines in current use will remain protective against the lambda variant and that monoclonal antibody therapy will remain effective.

## Introduction

The continued emergence of severe acute respiratory syndrome coronavirus 2 (SARS-CoV-2) variants with increased transmissibility poses concerns with regards to reinfection and diminished vaccine protection. The spread of variants also raises concerns regarding potential decrease in the efficacy of anti-spike protein monoclonal antibody therapy that has been shown to reduce disease symptoms and the rate of hospitalization^1^. Variants B.1.351 (Beta), B.1.617.2 (Delta), B.1.427/B.1.429 (Epsilon), B.1.526 (Iota), and B.1.1.248 (Gamma) encode spike proteins with L452R, E484K, E484Q mutations in the spike protein receptor binding domain (RBD) that provide a degree of resistance to neutralization by serum antibodies of vaccinated and convalescent individuals^2–7^.

The lambda variant is prevalent in Peru and is increasing in prevalence in neighboring Argentina, Ecuador, Chile and Brazil^8^. In June, the World Health Organization designated the variant (C.37 lineage) a variant of interest^9^. The variant spike protein is characterized by a novel deletion and mutations (Δ246-252, G75V, T76I, L452Q, F490S, T859N), L452Q and F490S of which are novel mutations in the RBD.

The increasing prevalence of the lambda variant raises concerns as to whether the current vaccines will contain its spread. In this study, we tested the sensitivity of viruses with the lambda variant spike protein to neutralization by convalescent sera, vaccine-elicited antibodies and Regeneron therapeutic monoclonal antibodies REGN10933 and REGN10987.

## Results

### Lambda spike protein-pseudotyped lentiviruses

The lambda spike protein has mutations L452Q and F490S in the RBD, and G75V, T76I mutations and 246-252 deletions in the N-terminal domain (NTD) **(Figure S1A)**. To analyze antibody neutralization of the variant spike protein, we generated expression vectors for the variant and its constituent mutations and used these to produce pseudotyped lentiviral virions encoding GFP and nano-luciferase reporters. The use of such pseudotypes to determine antibody neutralizing titers has been shown to yield results consistent with those obtained with the live virus plaque reduction neutralization test^10^. Immunoblot analysis of transfected pseudotype virus producer cells and virus-containing supernatants showed that the variant spike proteins were well expressed, proteolytically processed and incorporated into lentiviral virions at a level similar to that of the parental D614G spike protein **(Figure S1B)**. Analysis of the infectivity of the pseudotyped viruses on ACE2.293T cells, normalized for particle number, showed that the lambda spike protein increased infectivity by 2-fold. The increase was due to the L452Q mutation; the other mutations (G75V-T76I, F490S, T859N and Δ246-252) had no significant effect on infectivity (**Figure S1C**).

### Neutralization of the lambda variants by convalescent sera and vaccine-elicited antibody

Analysis of serum specimens from convalescent patients who had been infected prior to the emergence of the variants showed that viruses with the lambda variant spike protein were 3.3-fold resistant to neutralization by convalescent sera as compared to neutralization of virus with the parental D614G spike, similar to the 4.9-fold resistance of the B.1.351 variant to neutralization **(Figure 1A)**.

**Figure 1.**
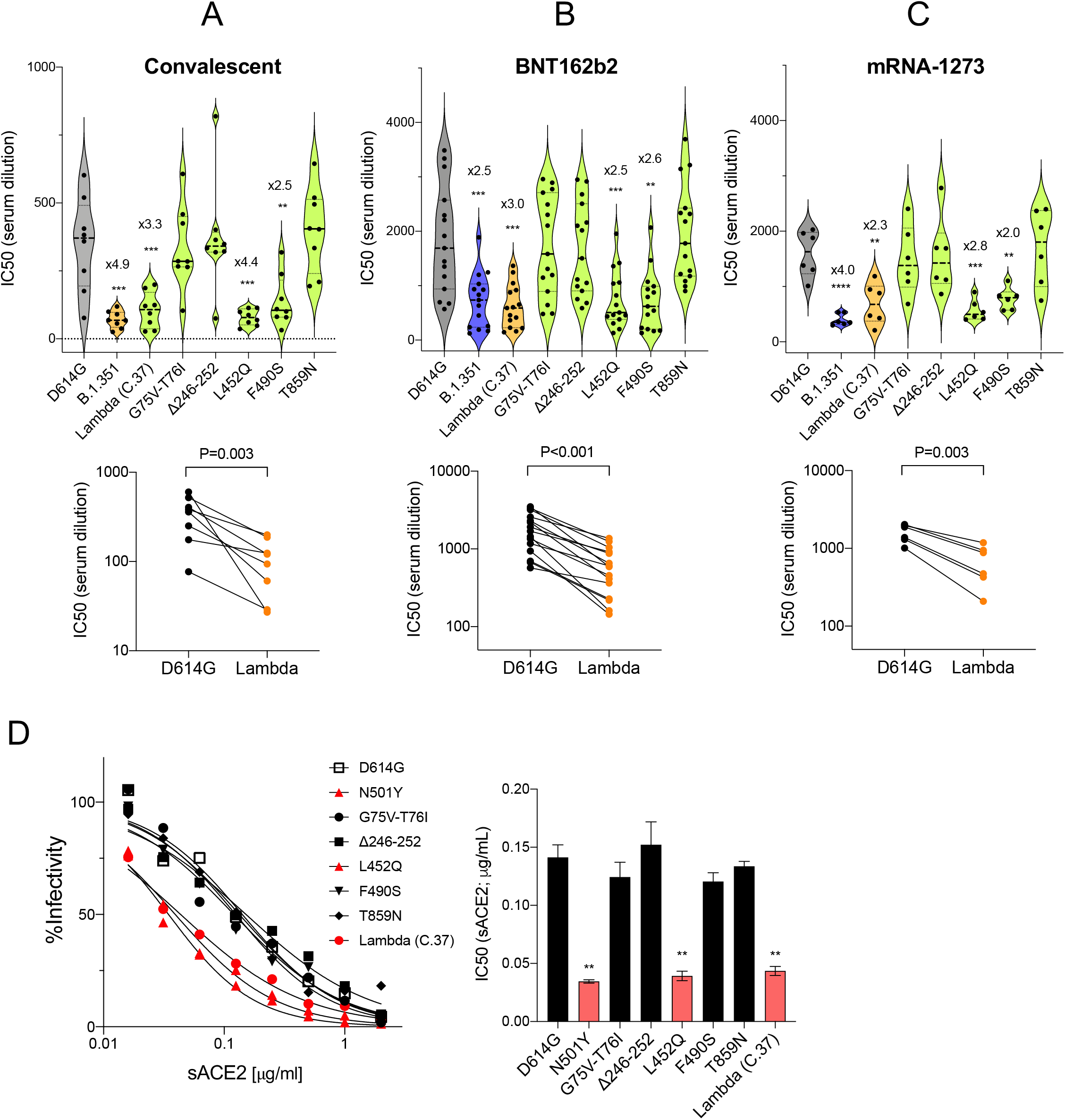
Neutralization of variant spike protein pseudotyped viruses by convalescent sera, vaccine-elicited antibodies, monoclonal antibodies and soluble ACE2. (A) Neutralization of lambda variant spike protein viruses pseudotyped virus by convalescent serum (n=8). Dots represent the IC50 of single donors. (B) Neutralizing titers of serum samples from BNT162b2 vaccinated individuals (n=15). Each dot represents the IC50 for a single donor. (C) Neutralizing titers of serum samples from mRNA-1273 vaccinated donors (n=6). The neutralization IC50 from individual donors is shown. Significance is based on the two-sided test. (**P≤0.05, ***P≤0.001, ****P≤0.0001). (D) Neutralization of beta (B.1.351) and lambda variant spike protein variants by REGN10933 and REGN10987 monoclonal antibodies. Neutralization of D614G and lambda variant pseudotyped viruses by REGN10933 (left), REGN10987 (middle), and 1:1 ratio of REGN10933 and REGN10987 (right). The IC50s of REGN10933, REGN10987 and the cocktail is shown in the table. (E) Neutralization of individual mutated spikes by REGN10933 (left), REGN10987 (middle), and cocktail (right). The table shows the IC50 of REGN10933, REGN10987 and the cocktail. (F) Neutralization of lambda variant spike protein variants by soluble sACE2. Viruses pseudotyped with variant spike proteins were incubated with a serially diluted recombinant sACE2 and then applied to ACE2.293T cells. Each plot represents the percent infectivity of D614G and other mutated spike pseudotyped virus. The diagram shows the IC50 for each curve.

Analysis of serum samples from individuals vaccinated with Pfizer BNT162b2 showed that virus with the lambda spike was about 3-fold resistant to neutralization **(Figure 1B).** Serum samples from individuals vaccinated with the Moderna mRNA-1273 vaccine were on average 2.3-fold resistant to neutralization **(Figure 1C).** The resistance was attributed to the L452Q and F490S mutations in the lambda spike protein **(Figure 1A, B, C).**

### L452Q increases spike protein affinity for ACE2

N501Y and L452R mutations in the RBD of earlier variants increase spike protein affinity for ACE2, an effect that most likely is a primary contributor to the increased transmissibility of the alpha, beta and delta variants^11^. To determine whether the lambda variant has an increased affinity for ACE2, we used a sACE2 neutralization assay in which pseudotyped virions were incubated with different concentrations of sACE2 and the infectivity of the treated virions was measured on ACE2.293T cells. The results showed that the lambda spike caused a 3-fold increase sACE2 binding. The increase was caused by the L452Q mutation and was similar to the increase provided by the N501Y mutation^12,13^ **(Figure 1D)**. The F490S mutation did not have a detectable effect on sACE2 binding. The findings suggest that L452Q, like L452R in the delta variant, increases virus affinity for ACE2, likely contributing to increased transmissibility.

### Neutralization by REGN10933 and REGN10987 monoclonal antibodies

Analysis of REGN10933 and REGN10987, the monoclonal antibodies that constitute the Regeneron REGN-COV2 therapy, showed that virus with the lambda variant spike protein was about 3.6-fold resistant to neutralization by REGN10987. The resistance was attributed to the L452Q mutation (Figure 2A and B). Virus with the lambda variant spike protein was neutralized by REGN10933 with no decrease in titer. The REGN10933 and REGN10987 cocktail neutralized the virus with no decrease in titer relative to virus with the D614G spike protein (Figure 2A and B).

**Figure 2.**
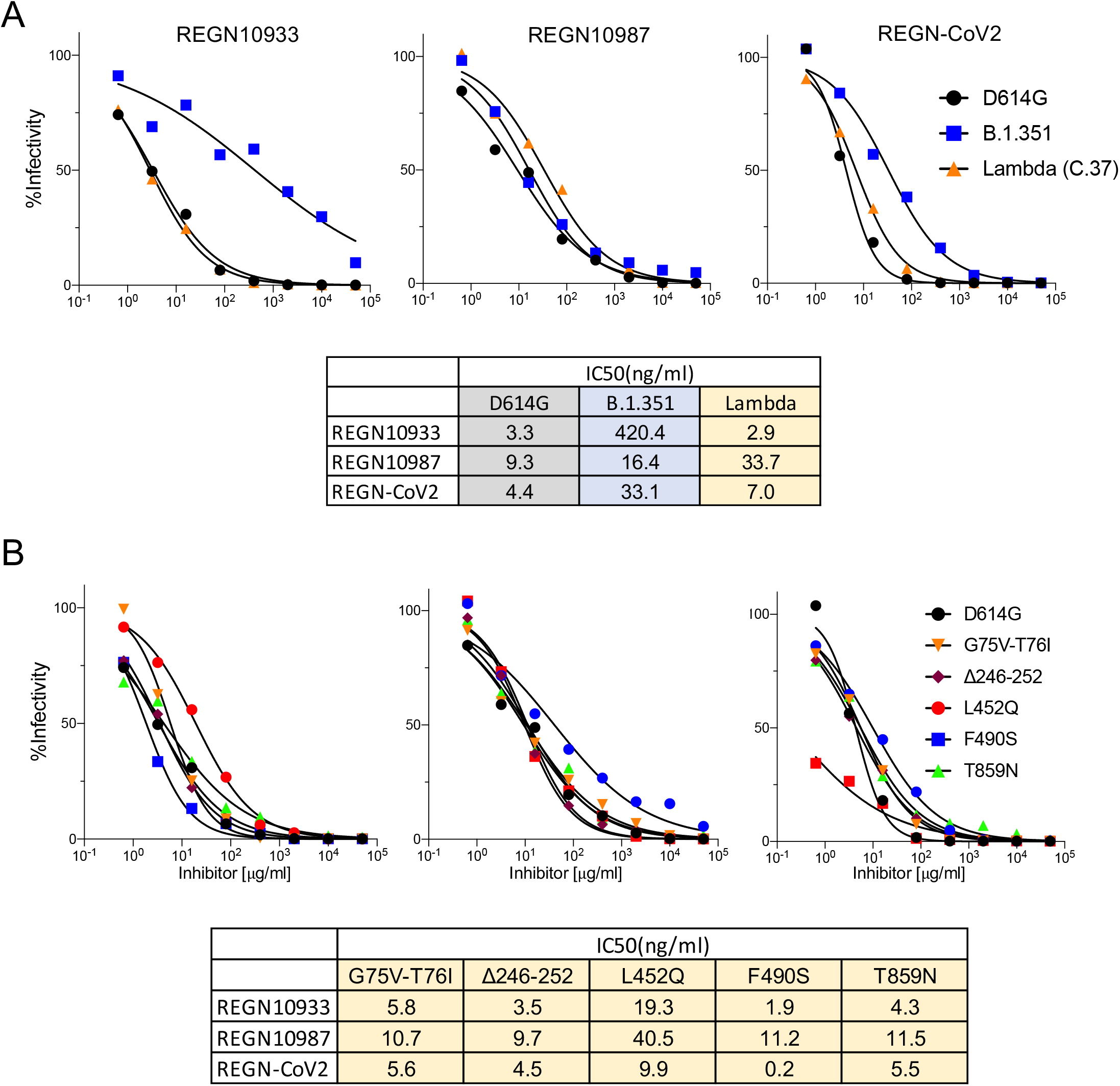

## Discussion

Virus with the lambda spike protein, like several VOC variant spike proteins showed a partial resistance to neutralization by vaccine-elicited antibodies and convalescent sera; however the average 3-fold decrease in neutralizing titer against the variant is not likely to cause a significant loss of protection against infection as the average neutralization IC50 titer by the sera of BNT162b2 and mRNA-1273 vaccinated individuals was about 1:600, a titer that is above that in the sera of individuals who recovered from infection with the parental D614G virus. A small fraction of vaccinated individuals had serum antibody titers less than average but whether this will lead to reduced protection from variant infection will need to be determined in epidemiological studies.

The resistance of the lambda variant to antibody neutralization was caused by the L452Q and F490S mutations. The L452R mutation of the California B.1.427/B.1.429 is associated with a 2-fold increase in virus shedding by infected individuals and a 4-6.7-fold and 2-fold decrease in neutralizing titer by the antibodies of convalescent and vaccinated donors, respectively^14^. The degree of neutralization resistance provided by L452Q was similar to that of L452R. Amino acid residues 490 and 484 lie close together on the top of the RBD and are therefore in a position to affect the binding of neutralizing antibody. The E484K mutation in the B.1.351, B.1.526, P.1 and P.3 spike proteins causes partial resistance to neutralization^2–7^. Similarly, the F490S mutation also caused a 2-3-fold resistance to neutralization, demonstrating the importance of the amino acid as an antibody recognition epitope. While the lambda variant was slightly resistant to REGN10987, it was neutralized well by the cocktail with REGN10933.

This study suggests that the L452Q and F490S mutations of the lambda variant spike protein caused a partial resistance to vaccine elicited serum and Regeneron monoclonal antibodies. While our findings suggest that current vaccines will provide protection against variants identified to date, the results do not preclude the possibility that novel variants will emerge that are more resistant to current vaccines. The findings highlight the importance of wide-spread adoption of vaccination which will protect individuals from disease, decrease virus spread and slow the emergence of novel variants.

## Acknowledgements

The work was funded by grants from the NIH to N.R.L. (DA046100, AI122390 and AI120898) and to M.J.M. (UM1AI148574), T.T. was supported by the Vilcek/Goldfarb Fellowship Endowment Fund.

## Author contributions

T.T. and N.R.L. designed the experiments. H.Z., T.T. and B.M.D. carried out the experiments and analyzed data. T.T., H.Z. and N.R.L. wrote the manuscript. M.I.S. and M.J.M. provided key reagents and useful insights. All authors provided critical comments on manuscript.

---

**Supplementary Figure 1.**
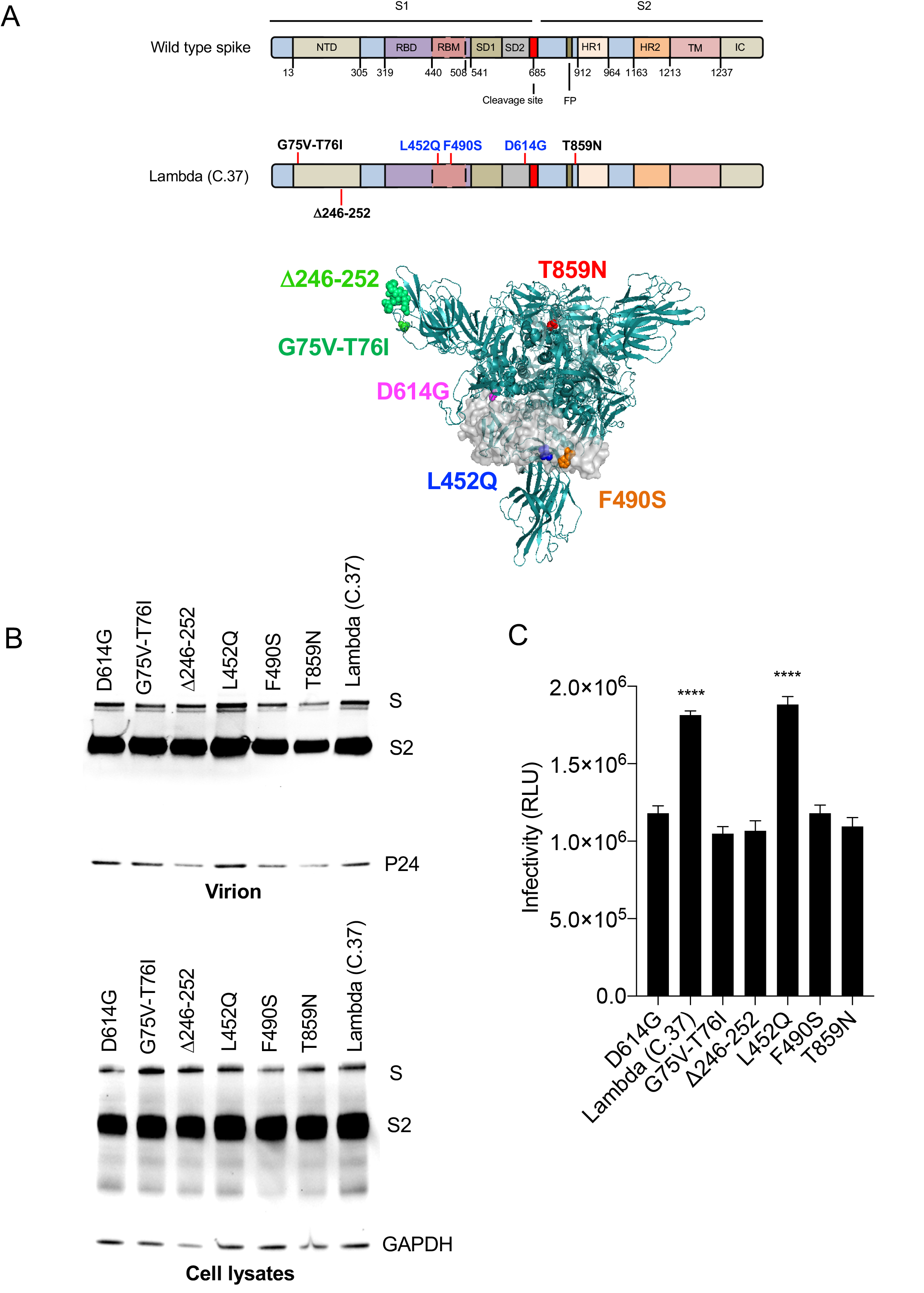
(A) The domain structure of the SARS-CoV-2 spike is diagrammed with the variant amino acid residues indicated. NTD, N-terminal domain; RBD, receptor-binding domain; RBM, receptor-binding motif; SD1 subdomain 1; SD2, subdomain 2; FP, fusion peptide; HR1, heptad repeat 1; HR2, heptad repeat 2; TM, transmembrane region; IC, intracellular domain. Key mutations are shown in 3D structure (top view). (B) Immunoblot analysis of the variant spike proteins in transfected 293T cells. Pseudotyped viruses were produced by transfection of 293T cells. Two days post-transfection, virions were analyzed on an immunoblot probed with anti-spike antibody and anti-HIV-1 p24. The cell lysates were probed with anti-spike antibody and anti-GAPDH antibodies as a loading control. (C) Infectivity of virus pseudotyped by lambda variant and D614G spike proteins. Viruses were normalized for RT activity and applied to target cells. Infectivity of viruses pseudotyped with the lambda variant protein or the individual lambda mutations were tested on ACE2.293T. Luciferase activity was measured two days post-infection. Significance was based on two-sided testing. (**P≤0.05, ***P≤0.001).

## Methods

### Plasmids

Mutations in the spike were introduced into pcCOV2.Δ19.D614GS by overlap extension PCR and confirmed by DNA sequencing. Plasmids used in the production of lentiviral pseudotyped virus have been previously described^2^.

### Human Sera and monoclonal antibodies

Convalescent sera and BNT162b2 or Moderna-vaccinated sera were collected on day 28, 7 days post-second immunization, at the NYU Vaccine Center with written consent under IRB approved protocols (IRB 18-02035 and IRB 18-02037). Donor age and gender were not reported. Regeneron monoclonal antibodies (REGN10933 and REGN10987) were generated as previously described^11^.

### SARS-CoV-2 spike lentiviral pseudotypes

Lentivirus pseudotyped by variant SARS-CoV-2 spikes were produced as previously reported^2^. Viruses were concentrated by ultracentrifugation and normalized for reverse transcriptase (RT) activity. To determine neutralizing activity, sera or monoclonal antibodies were serially diluted and then incubated with pseudotyped virus (approximately 2.5 × 10^7^ cps) for 30 minutes at room temperature and then added to target cells. Luciferase activity was measured 2 days post infection^2^.

### Soluble ACE2 Neutralization assay

Serially diluted recombinant soluble ACE2 protein prepared from transfected CHO cells was incubated with pseudotyped virus for 30 minutes at room temperature and added to ACE2.293T cells. Luciferase activity was measured using Nano-Glo luciferase substrate (Nanolight) in an Envision 2103 microplate luminometer (PerkinElmer).

### Immunoblot analysis

Proteins were analyzed on immunoblots probed with mouse anti-spike monoclonal antibody (1A9) (GeneTex), anti-p24 monoclonal antibody (AG3.0) and anti-GAPDH monoclonal antibody (Life Technologies) followed by goat anti-mouse HRP-conjugated secondary antibody (Sigma) as previously described^2^.

### Statistical Analysis

All experiments were in technical duplicates or triplicates and the data were analyzed using GraphPad Prism 8. Statistical significance was determined by the two-tailed, unpaired t-test. Significance was based on two-sided testing and attributed to p< 0.05. Confidence intervals are shown as the mean ± SD or SEM. (*P≤0.05, **P≤0.01, ***P≤0.001, ****P≤0.0001). The PDB file of SARS-CoV-2 spike protein (7BNM)^15^ was downloaded from the Protein Data Bank. 3D view of protein was obtained using PyMOL.

